# Quantifying the Payments for Ecosystem Services among hydrologic units in Zhujiang River Basin, China based on the indicator of Optional Capacity Value

**DOI:** 10.1101/616680

**Authors:** Haile Yang, Bin Zhao, Jiakuan Chen

## Abstract

Ecosystem services (ES) are fundamental to human being’s livelihoods, production and survival. However, the spatial mismatch between ES supply and demand is a common phenomenon. Payments for Ecosystem Services (PES) provide a way to promote the complementary advantages and benefits equilibrium between ES supplier and beneficiary. At present, PES is mainly based on the tradeoff between the profit and loss of ecological conservation. The quantifying of PES mainly uses the opportunity cost of ES supplier and follows the principle of additionality, which neglects the benefits that arise from the basic (contrast to additional) ES experienced by ES beneficiary and ignores the rights and interests of ES supplier who supplies the basic ES. To resolve this problem, we proposed that we should set the value of ES experienced by ES beneficiary as the quantitative indicator of PES. Here, we introduced a new indicator (optional capacity value, OCV) to implement this idea. The ES OCV indicates the optional capacity of supporting the total value produced by human being’s economic and social activities provided by the total volume of an ES. In this paper, we calculated the ES OCV of water provision in Zhujiang River Basin (Pearl River Basin), China. Then, we discussed three scenarios of quantifying PES, based on the principles of (1) interests sharing and responsibilities bearing and (2) equal pay for equal work. The results showed that the ES OCV could describe the conditions that water resources in a hydrologic unit not only provide benefits to the hydrologic unit itself, but also provide benefits to downstream hydrologic units, and then could be a quantitative indicator for PES. This research provides a new PES scheme which would promote the coordinated development and ecological conservation among the regions with mismatch between ES supply and demand.

## 1 Introduction

Ecosystem services (ES) are the benefits that humans derive from nature and are fundamentally important for human wellbeing, health, livelihoods, and survival (Costanza et al., 1997; Millennium Ecosystem Assessment, 2005; TEEB Synthesis, 2010; Mancini et al., 2018). Mismatches between ES supply and demand are common in the world (Geijzendorffer et al., 2015; Baro et al., 2016; Wei et al., 2017). Payments for Ecosystem Services (PES) provide a pathway that ES beneficiary pays something to ES supplier as compensation, which reflects the benefits experienced by ES beneficiary and the costs paid out by ES supplier (Engel et al., 2008; Zheng et al., 2013; Wunder, 2015). PES promotes the optimization of landscape planning and land use management, the complementary advantages and benefits equilibrium between ES supplier and beneficiary, and then increases the regional and global ES provision (Farley and Costanza, 2010; Hayha and Franzese, 2014; Fu et al., 2018). We could qualitatively or quantitatively describe the value of ES (Costanza et al., 1997; Mancini et al., 2018; Martin and Mazzotta, 2018). The ES value has been used to support ecosystem management (Grizzetti et al., 2016; Kubiszewski et al., 2017; Sun et al., 2018) and is expected to provide a quantitative indicator for PES.

At present, PES is mainly based on the tradeoff between the profit/loss of ES supplier and beneficiary in natural resources exploitation or in ecological conservation (Engel et al., 2008) and follows the principle of additionality which means that ES beneficiary only pays for the ecological conservation which could produce additional ES (Wunder, 2015). This PES approach neglects the benefits arising from the basic (contrast to additional) ES experienced by ES beneficiary and ignores the rights and interests of ES supplier who supplies the basic ES. To resolve this problem, we need to quantify PES based on the benefits (ES values) experienced by ES benefic iary. However, current ES valuation approaches could not provide any indicator to quantify the PES based on the basic ES. So, we need a new approach and a new indicator to value ES.

PES could be user-financed, in which funding comes from the users of the ES, or be government-financed, in which funding comes from a third party (typically the government) (Engel et al., 2008; Wunder et al., 2008; Wunder, 2015). Theoretically, user-financed PES is more effective and flexible than government-financed PES in a free market (Pagiola and Platais, 2006). Practically, government-financed PES is more cost-efficient than user-financed PES because of transaction cost, especially when the PES program is conducted in a large-scale (Engel et al., 2008). Moreover, most of the PES programs in both developed and developing countries are government-financed (Schomers and Matzdorf, 2013). So, we expect the new ES value indicator should be adapted to government-financed PES.

As we all know, human beings could not survive without any of vital ES which is indispensable to human being’s survival (such as oxygen, freshwater) (Chaisson, 2002). In another word, all values produced by human depend on vital ES. In another perspective, ES could not provide any values to people without the presence of human capital, social capital and built capital (Costanza et al., 2014). So, the consumption of vital ES supports the total value produced by human being’s economic and social activities (TVPH). The total volume of ES provides the freedom of choice for ES consumption. So, the ES value provided by a vital ES could be described as the optional capacity for supporting the TVPH, which could be calculated by using the product of multiplying the TVPH by the freedom of choosing the consumption from the total volume of this ES. This new ES value is termed as ES optional capacity value (OCV).

In this work, we presented a case study of valuing the ES OCV of water provision in Zhujiang River Basin (Pearl River Basin), China to quantify PES. The main objectives of this study are: (1) to discuss how the ES OCV serves as a quantitative indicator for PES; (2) to show the approach of valuing ES OCV. This study would provide a new scheme and tool for understanding and implementing PES.

## 2 Materials and methods

### 2.1 Study area and data

The Zhujiang River Basin in South China covers an area of 453,700 square km and consists of four major river systems, the Xijiang River, the Beijiang River, the Dongjiang River and the Zhujiang Delta (ECERLC, 2013). In the Water Resources Bulletin published by the Pearl River Water Resources Commission of the Ministry of Water Resources (PRWRC), the Zhujiang River Basin is divided into 7 hydrologic units, the Nanpan-Beipan River, the Hong-Liu River, the Yujiang River, the Xijiang River Lower Reach, the Beijiang River, the Dongjiang River and the Zhujiang Delta (Fig. 1) (PRWRC, 2017).

**Fig. 1.**
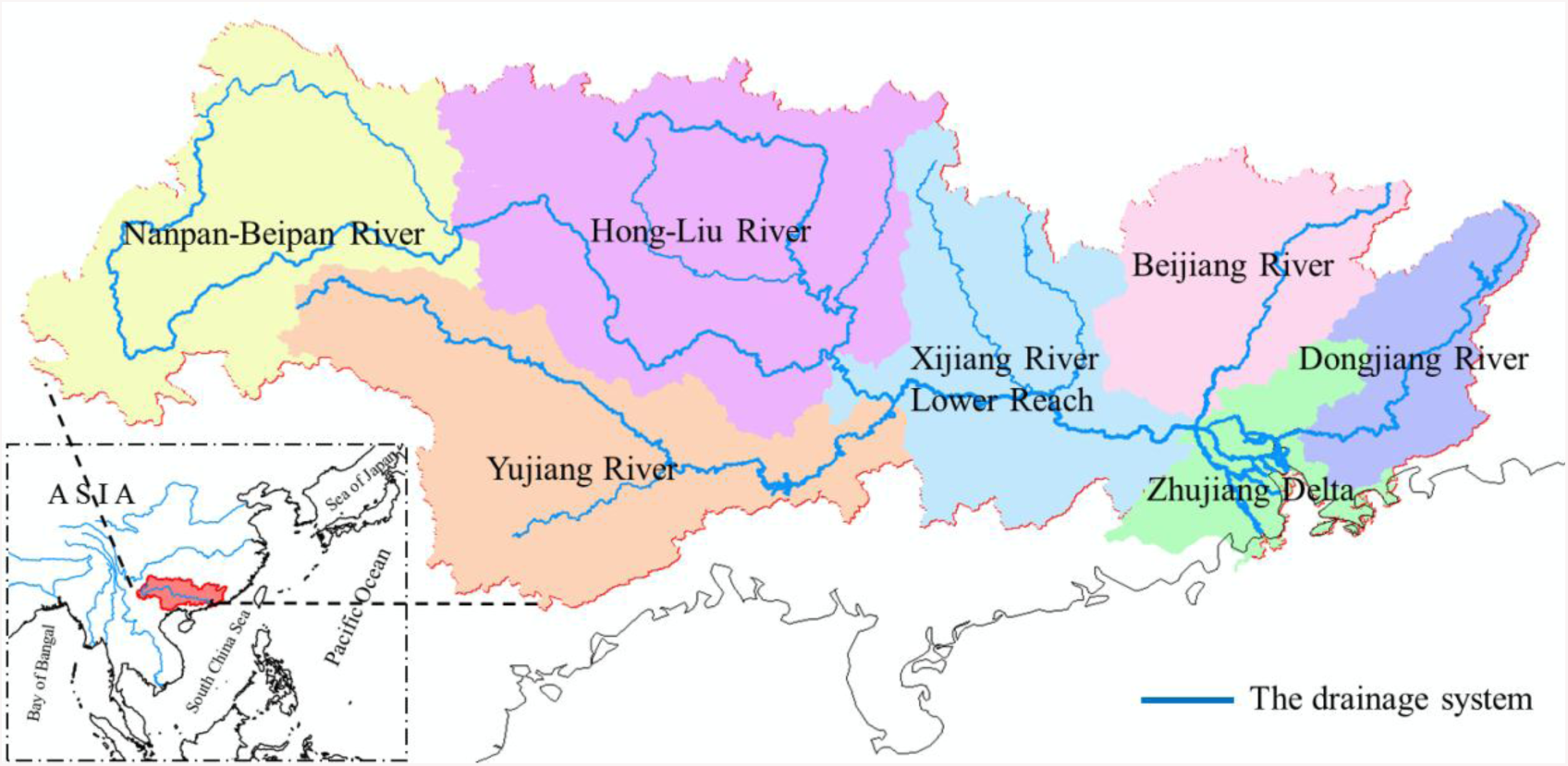
Zhujiang River Basin (Pearl River Basin) and its seven hydrologic units

Water provision is a vital ES for human being’s survival and a most widely cited ES. In many studies on water provision, the quantity, quality and spatial-temporal distribution of water resources are analyzed separately (Hackbart et al., 2017). In this work, we just used the water quantity for evaluating the ES OCV of water provision, and then quantified the PES among the hydrologic units of Zhujiang River Basin. All data (i.e. water resources, water consumption and gross domestic product in Zhujiang River Basin and its seven hydrologic units) used in this work comes from the Water Resources Bulletin published by PRWRC (http://www.pearlwater.gov.cn/xxcx/szygg/) (PRWRC, 2017).

### 2.2 Methods

The ES value of a vital ES which is indispensable for human being’s survival is the optional capacity for supporting the TVPH, which could be calculated by using the product of multiplying the TVPH by the freedom of consumption. In this work, we used the average uncertainty of choosing the ES consumption from its total volume to indicate the freedom of consumption, in which the average uncertainty was described by log base 2 which indicated the uncertainty in a binary decision and was valued in bits (Ulanowicz, 1986). In calculating the OCV of ES spatial subsidy (the ES comes from other regions), we followed the principle of local priority (Eq. 1, 2).

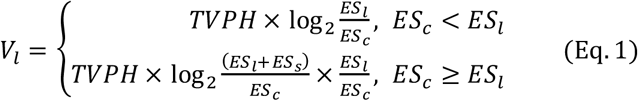

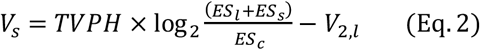

In the equations, *V*_*l*_ denotes the local ES OCV in a region; *V*_*s*_ denotes the spatial subsidized ES OCV in a region; *ES*_*c*_ denotes the volume of ecosystem service consumed in a region, TVPH denotes the total value produced by human being’s economic and social activities.

In this case study, we set the 7 hydrologic units of Zhujiang River Basin as the accounting units, used the gross domestic product (GDP) to indicate the TVPH, water consumption to indicate ES consumption, local water yield to indicate local ES, passing-by water to indicate ES spatial subsidy, and then (1) calculated the ES OCV of water provision experienced by each hydrologic unit which included the ES OCV of local water provision and of input water provision, (2) calculated the ES OCV of water provision provided by each hydrologic unit which included the ES OCV of local water provision and of output water provision.

As a hydrologic unit not only provides ES OCV of water provision to itself but also to downstream hydrologic units, PES should follow the principles of (1) interests sharing and responsibilities bearing and (2) equal pay for equal work. The principle of interests sharing and responsibilities bearing means that (1) all hydrologic units that experience the ES of water provision supplied by hydrologic unit A should pay to hydrologic unit A, and (2) the payments should be positively related to their ES OCV. The principle of equal pay for equal work means that (1) hydrologic unit B should pay to all hydrologic units which supply the ES of water provision to hydrologic unit B, and (2) the payments should be positively related to their ES OCV.

## 3 Results

### 3.1 ES OCVs of water provision provided and experienced by each hydrologic unit of Zhujiang River Basin

According to calculate the ES OCV of water provision experienced by each hydrologic unit of Zhujiang River Basin in 2016, we found that the Hong-Liu River, Xijiang River Lower Reach and Zhujiang Delta benefited from both local water yield and passing-by water. In 2016, the Hong-Liu River, Xijiang River Lower Reach and Zhujiang Delta respectively received passing-by water 26.95 billion m^3^, 161.16 billion m^3^ and 347.35 billion m^3^ which provided ES OCV 0.1839 trillion yuan bits, 0.9584 trillion yuan bits and 18.4487 trillion yuan bits, respectively (Table 1).

**Table 1.** The ES OCV (*V*) of water provision (including local water yield and passing-by water) experienced by each hydrologic unit of Zhujiang River Basin in 2016

|  | Total volume<br>(e9 m<sup>3</sup>) | Usage (e9<br>m<sup>3</sup>) | GDP (e12<br>yuan) | V (e12<br>yuan bits) |
| --- | --- | --- | --- | --- |
| Nanpan-Beipan<br>River | 29.56 | 4.78 | 0.6373 | 1.6753 |
| Hong-Liu River | 105.98 | 9.17 | 0.5626 | 1.9863 |
| Passing-by water | 26.95 |  |  | 0.1839 |
| Yujiang River | 35.70 | 7.87 | 0.5745 | 1.2532 |
| Xijiang River Lower<br>Reach | 83.48 | 10.38 | 0.6179 | 1.8583 |
| Passing-by water | 161.16 |  |  | 0.9584 |
| Beijiang River | 69.64 | 5.26 | 0.4109 | 1.5315 |
| Dongjiang River | 41.95 | 4.51 | 1.0488 | 3.3746 |
| Zhujiang Delta | 37.93 | 17.10 | 5.5161 | 6.3399 |
| Passing-by water | 347.35 |  |  | 18.4487 |
| **Sum** | 939.70 | 59.07 | 9.3681 | 37.6100 |

Distributing the ES OCV of passing-by water into the hydrologic units that export water resources, we got the total ES OCV of water provision provided by each hydrologic unit, including local services and exported services (Table 2). Among 7 hydrologic units of Zhujiang River Basin, there were a series of ES OCV spatial subsidies (Table 2).

**Table 2.**
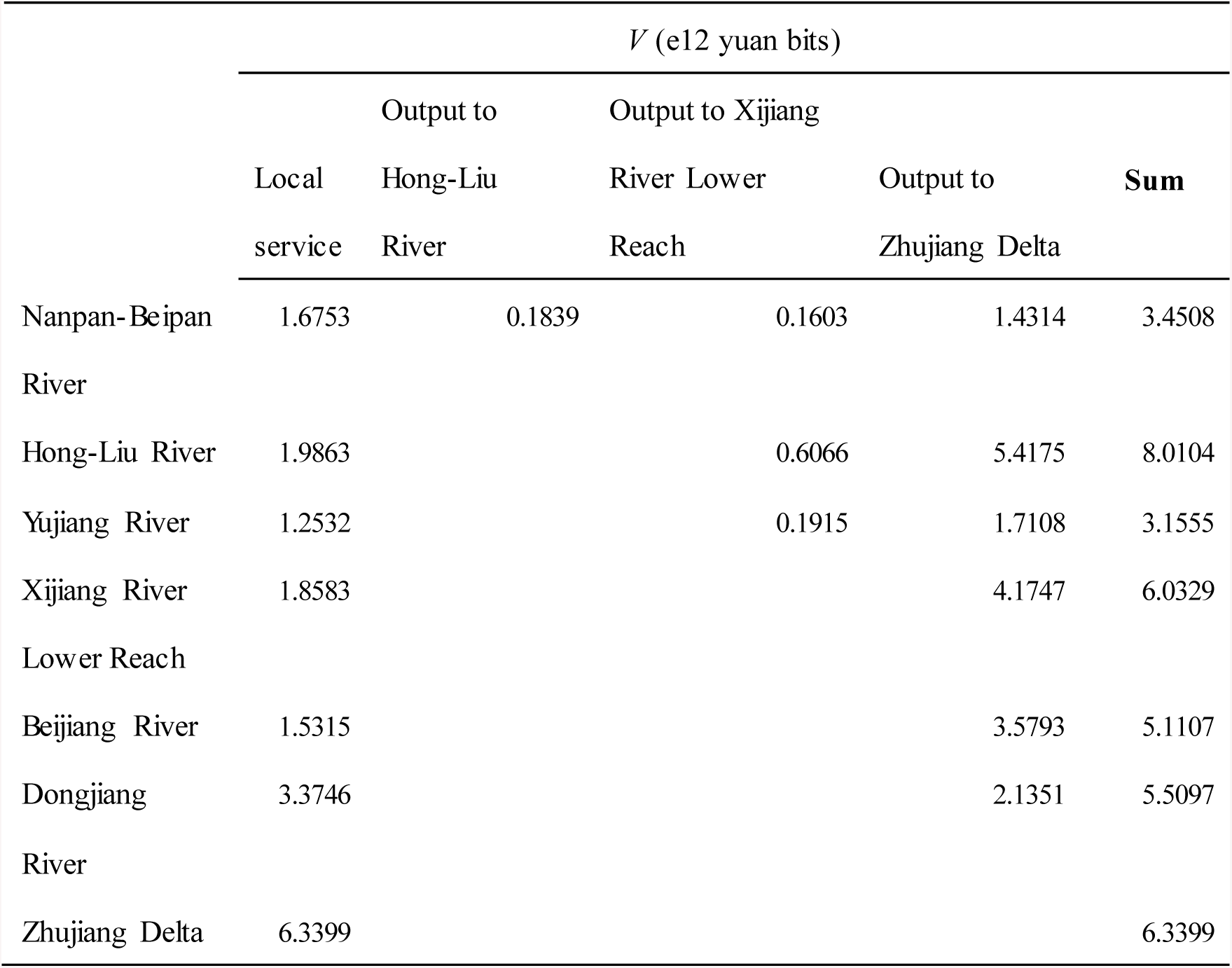
The ES OCV (*V*) of water provision (including local yield and passing-by water) provided by each hydrologic unit of Zhujiang River Basin in 2016

### 3.2 PES among hydrologic units of Zhujiang River Basin based on the ES OCVs of water provision

The ES OCVs of water provision provided and experienced by each hydrologic unit of Zhujiang River Basin provide quantitative indicators for PES among hydrologic units. Following the principle of interests sharing and responsibilities bearing, if there is a water resources protection project in the hydrologic unit of Nanpan-Beipan River with a budget of 3.4508 billion yuan, then the hydrologic units experienced the ES of water provision supplied by the hydrologic unit of Nanpan-Beipan River should apportion this cost based on their ES OCVs (Fig. 2).

**Fig. 2.**
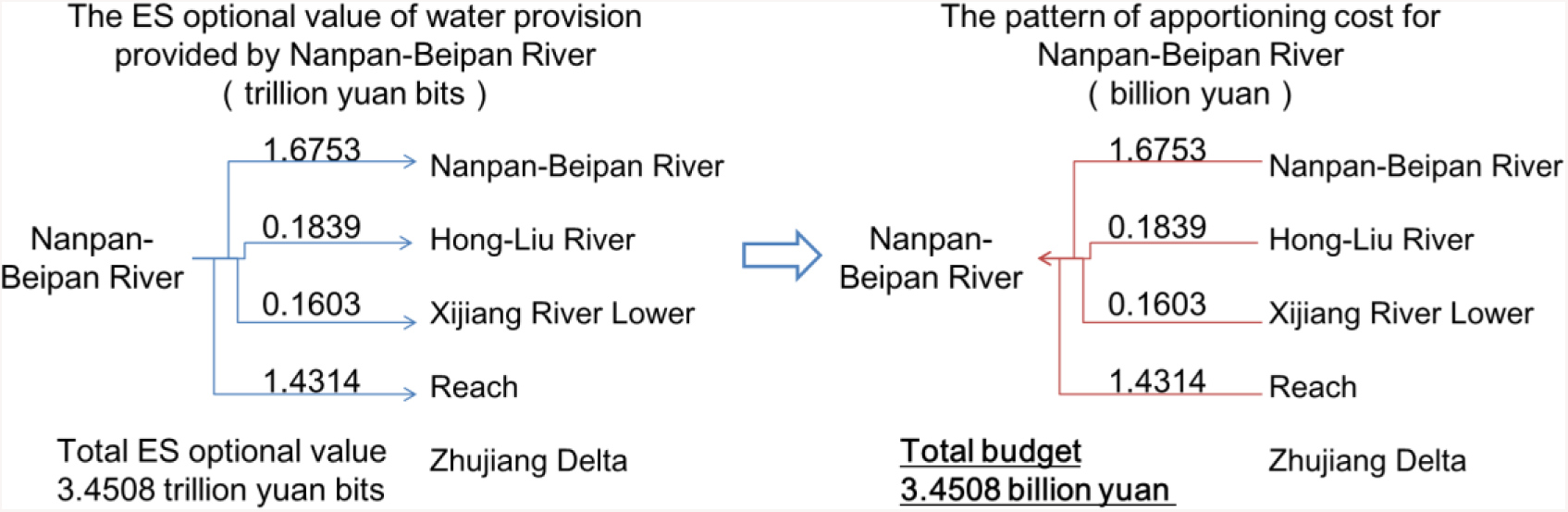
Pattern of apportioning cost based on the ES OCV in 2016. Here we assumed that the water resource protection project in Nanpan-Beipan River had a budget of 3.4508 billion yuan.

Following the principle of equal pay for equal work, if there is a water resources protection project in the hydrologic unit of Zhujiang Delta with a budget of 6.3399 billion yuan, then the hydrologic unit of Zhujiang Delta should pay for the ES of passing-by water to the hydrologic units of exporting water resources to it based on their ES OCVs (Fig. 3).

**Fig. 3.**
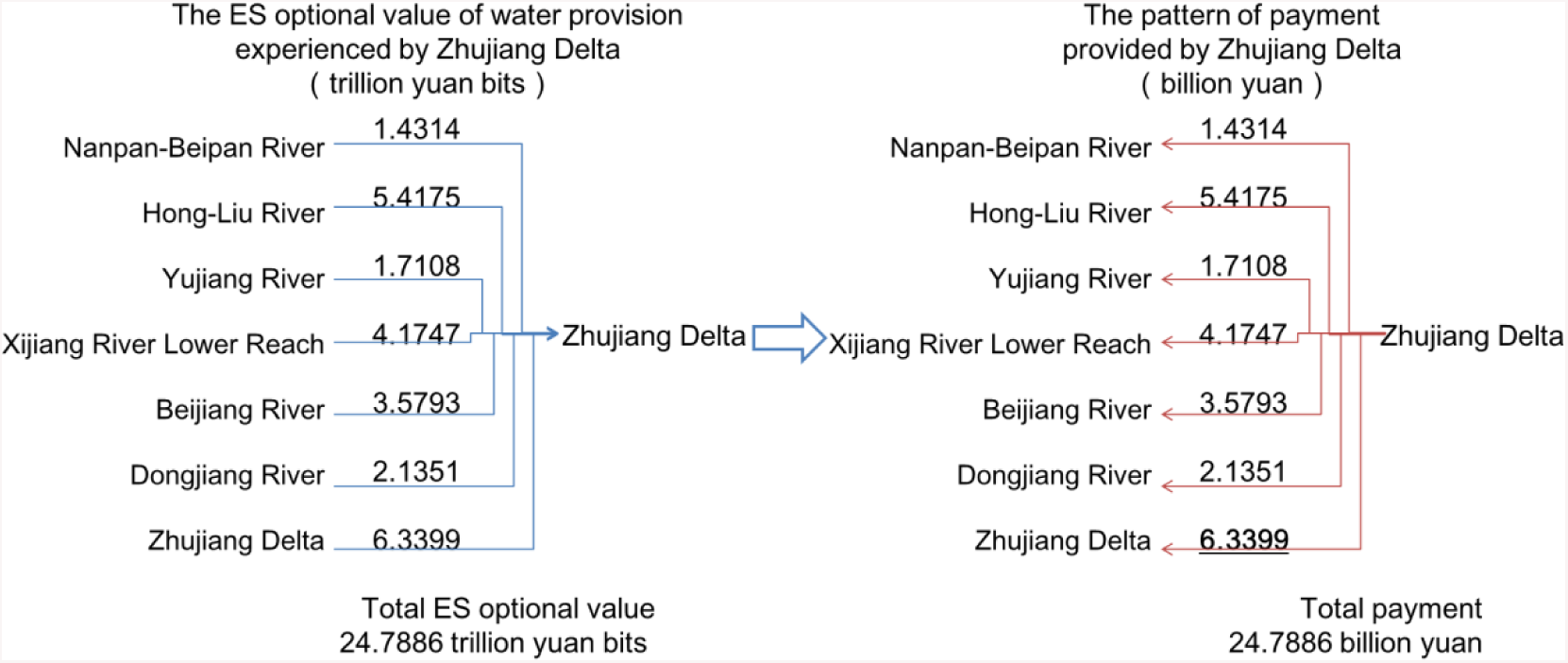
Pattern of payment for water resources protection based on the ES OCV in 2016. Here we assumed that the water resources protection project in Zhujiang Delta had a budget of 6.3399 billion yuan.

Combining the principle of interests sharing and responsibilities bearing and the principle of equal pay for equal work, if there is a water resource protection project at the Zhujiang River Basin scale, then the project funding should be apportioned among all hydrologic units based on the ES values of water provision experienced by each hydrologic unit, and be distributed to each hydrologic unit based on the ES values of water provision provided by each hydrologic unit (Fig. 4).

**Fig. 4.**
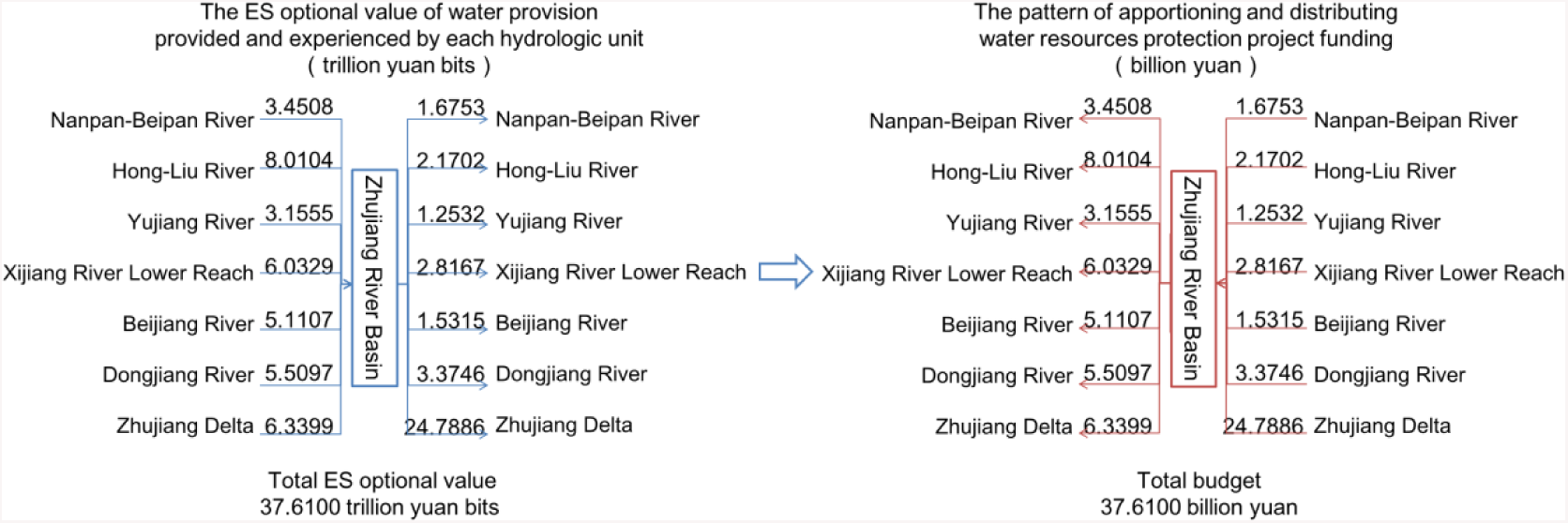
Pattern of apportioning and distributing the project funding for the water resources protection project in Zhujiang River Basin based on the ES value in 2016. Here we assumed that its total budget was 37.6100 billion yuan.

## 4 Discussion

### 4.1 Quantitative indicator for PES

We need a new PES scheme that quantifies PES based on the ES value experienced by beneficiary. Traditional PES approaches are based on the tradeoff between the profit/loss of ES supplier and beneficiary in natural resources exploitation or in ecological conservation (Engel et al., 2008) and use the ES increment as the reference for PES (Wunder, 2015). That ignores the rights and interests configuration of basic ES between supplier and beneficiary. The same as mineral resources, energy resources, land resources, channel resources and so on, ES has its owner (i.e. supplier) and should get the compensation provided by ES beneficiary. As it cannot control the ES supply sufficiently, the ES supplier always is the weak in the supply-demand relationship and seldom gets an adequate compensation. To preserve the rights of ES supplier and quantify the benefits of basic ES experienced by beneficiary and, we need to quantify PES based on the ES value experienced by beneficiary.

ES OCV provides a quantitative indicator for the new PES scheme. To implement the new PES scheme – quantifying PES based on the ES value experienced by beneficiary, we need to appropriately quantify its ES value, firstly. Traditional monetary ES valuation approaches are constrained by the complexities of ecosystem and the limitations of economic methodology, which result in too large (Costanza et al., 2014; Dai et al., 2016) and too diverse (Costanza et al., 2014; Kubiszewski et al., 2017) ES values to use in practical PES (Liu et al., 2018). The results of traditional non-monetary ES valuation always are a physical index or a set of physical indexes (Mancini et al., 2018; Martin and Mazzotta, 2018), which could not reflect the benefits arising from ES experienced by beneficiary. ES OCV reflects the level of economic development, the ES consumption and the total ES supply (Eq. 1, 2). So, it could quantify the ES value of local services and imported services experienced by a region (Table 1, 2), which provides a quantitative indicator for quantifying PES (Fig. 2, 3, 4). At the same time, it quantifies the ES value of local services and exported services provided by a region (Table 1, 2), which provides a quantitative reference for quantifying PES (Fig. 2, 3, 4). In the new PES scheme, we set regions as accounting units, set ES OCV fluxes among accounting units as indicators for PES, follow the principles of (1) interests sharing and responsibilities bearing and (2) equal pay for equal work, quantify the PES for basic ES (Fig. 2, 3, 4), and then provide quantitative indicators for government-financed PES. Moreover, comparing with the traditional valuation approaches which could provide a definite compensation for PES (Fu et al., 2018), our new PES scheme is more transparent, effective and flexible in practice.

The new PES scheme based on ES OCV provides a new pathway for the PES in Zhujiang River Basin, and could provide a reference for promoting the coordinated development and ecological conservation in other regions. Zhujiang Delta is one of the most crowded and developed areas in China. However, the social and economic development in Zhujiang River Basin is unbalanced (PRWRC, 2017). In 2014, the State Council of the People’s Republic of China approved the Development Planning of Zhujiang-Xijiang Economic Zone, which emphasized the coordinated regional development and the watershed ecological civilization construction (SCPRC, 2014). Constructing an appropriate basin ecological compensation approach is an important work. The exploration of this paper provided a new pathway for basin ecological compensation. We valued the ES OCV of water resources fluxes from upstream hydrologic units to downstream hydrologic units of Zhujiang River Basin as an indicator for PES, valued the ES OCV of water provision provided and experienced by each hydrologic unit as a reference frame, and then followed definite principles to quantify PES. The principle of “interests sharing and responsibilities bearing” means that all ES beneficiaries should pay to the ES supplier using the basic ES OCVs experienced by each hydrologic unit as quantitative indicators (Fig. 2, 4), which achieves the equity of PES among hydrologic units at basic ES level. The principle of “equal pay for equal work” means that a downstream hydrologic unit should pay to upstream hydrologic units, if it pays for local water resources protection program, using the basic ES OCVs provided by each hydrologic unit as quantitative indicators (Fig. 3, 4), which achieves the equity of PES among ES OCVs at basic ES level. The new PES scheme could promote the ecological conservation of upstream, the ES usage of downstream and the regional coordinated development. Similar to Zhujiang River Basin, the unbalanced social-economic development and the mismatch between ES supply and demand are common in the world (Morri et al., 2014; Wei et al., 2017). Our new PES scheme could be used to deal with these mismatches too.

### 4.2 Approaches of ES valuation

As water provision is a vital ES for human being’s survival, we described its ES OCV as the optional capacity for supporting the TVPH, calculated its ES OCV by using the product of multiplying the TVPH by the freedom of consumption (Eq. 1, 2). To keep the statistical caliber to be consistent, we completely used the data sets come from the Water Resources Bulletin published by PRWRC. In the Water Resources Bulletin, GDP is the most appropriate indicator to indicate the TVPH in all social and economic indexes, so we used GDP to indicate the TVPH in this case study (Table 1). In future, if there is another indicator which could indicate the TVPH more accurately than GDP, we could use that indicator to replace GDP. Of course, this replacement has no impact on the assessment scheme of ES OCV.

Here, the freedom was evaluated by the average uncertainty of selecting ES consumption from the total volume of this ES. The average uncertainty was described by log base 2 which indicated the uncertainty in a binary decision (Ulanowicz, 1986). If someone believes that another indicator or method could indicate or calculate the average uncertainty more appropriately, he could use that one to replace log base 2 in his study. Of course, this replacement does not change the assessment scheme of ES OCV.

In this work, the index of ES OCV and its valuation approach are different from traditional ES value indexes and valuation approaches. Traditionally, the valuation of ES could be monetary, using monetary unit to quantify ES value (Guo et al., 2001; Costanza et al., 2014; Dai et al., 2016; Kubiszewski et al., 2017); could be non-monetary, using biophysical indexes (Coscieme et al., 2014; Mancini et al., 2018), or a set of biophysical, sociocultural, economic, health and holistic indexes (Pascual et al., 2017; Folkersen, 2018; Martin and Mazzotta, 2018) to quantify ES value; could be mixes of monetary and non-monetary (Hayha et al., 2015; Kenter et al., 2016). These traditional ES valuation approaches focus on the value of ES in one region or the value of ES flow among regions, which are based on static standards arising from a definite area and a definite time. The ES OCV in this work indicates the optional capacity for supporting the TVPH, which depends on the level of economic development, the ES consumption and the total ES supply of the region that experience ES (Eq. 1, 2). It could dynamically reflect the ES value experienced by a region, including local services and imported services, and the ES value provided by a region, including local services and exported services (Table 1).

## 5 Conclusion

In this work, we analyzed the ES OCVs of water provision provided and experienced by 7 hydrologic units of Zhujiang River Basin, China, and the ES OCV fluxes among these hydrologic units. Then, we discussed the PES among 7 hydrologic units based on these ES OCVs and ES OCV fluxes. A hydrologic unit not only provides ES value of water provision to itself but also to downstream hydrologic units. Local water yield in a hydrologic unit provides ES OCV to this hydrologic unit, so this hydrologic unit should bear the responsibility of water resources protection. Passing-by water provides ES OCV to recipient hydrologic unit, so recipient hydrologic unit should pay to supplier hydrologic units. Following the principles of (1) interests sharing and responsibilities bearing and (2) equal pay for equal work, the ES OCVs of water provision provided and experienced by hydrologic units provide quantitative indicators for PES among hydrologic units. This work proposed a new method to quantify PES, which would promote the regional coordinated development and ecological conservation.

ES: ecosystem service(s)
OCV: optional capacity value
GDP: gross domestic product
PES: payments for ecosystem services
TVPH: total value produced by human being’s economic and social activities
PRWRC: Pearl River Water Resources Commission of the Ministry of Water Resources

## Acknowledgments

We thank Leon C Braat, Li Yongchi, and Li Zhaolei for grateful assistance that helps to improve the manuscript. This research did not receive any specific grant from funding agencies in the public, commercial, or not-for-profit sectors.

## References

Baro, F., Palomo, I., Zulian, G., Vizcaino, P., Haase, D. and Gomez-Baggethun, E., 2016. Mapping ecosystemservice capacity, flow and demand for landscape and urban planning: A case study in the Barcelona metropolitan region. LAND USE POLICY, 57:405–417. https://doi.org/10.1016/j.landusepol.2016.06.006

Chaisson, E., 2002. Cosmic Evolution: The Rise of Complexity in Nature. Harvard University Press, Cambridge, Massachusetts.

Coscieme, L., Pulselli, F.M., Marchettini, N., Sutton, P.C., Anderson, S. and Sweeney, S., 2014. Emergy and ecosystemservices: A national biogeographical assessment. ECOSYST SERV, 7:152–159. https://doi.org/10.1016/j.ecoser.2013.11.003

Costanza, R., DArge, R., DeGroot, R., Farber, S., Grasso, M., Hannon, B., Limburg, K., Naeem, S., ONeill, R.V., Paruelo, J., Raskin, R.G., Sutton, P. and VandenBelt, M., 1997. The value of the world’s ecosystemservices and natural capital. NATURE, 387:253–260. https://doi.org/10.1038/387253a0

Costanza, R., de Groot, R., Sutton, P., van der Ploeg, S., Anderson, S.J., Kubiszewski, I., Farber, S. and Turner, R.K., 2014. Changes in the global value of ecosystemservices. GLOBAL ENVIRON CHANG, 26:152–158. https://doi.org/10.1016/j.gloenvcha.2014.04.002

Dai, Z., Puyang, X. and Han, L., 2016. Using assessment of net ecosystemservices to promote sustainability of golf course in China. ECOL INDIC, 63:165–171. https://doi.org/10.1016/j.ecolind.2015.11.056

ECERLC: Editorial Committee of Encyclopedia of Rivers and Lakes in China, 2013. Encyclopedia of Rivers and Lakes in China, Section of Zhujiang River Basins. China Water & Power Press, Beijing (in Chinese).

Engel, S., Pagiola, S. and Wunder, S., 2008. Designing payments for environmental services in theory and practice: An overview of the issues. ECOL ECON, 65:663–674. https://doi.org/10.1016/j.ecolecon.2008.03.011

Farley, J. and Costanza, R., 2010. Payments for ecosystemservices: From local to global. ECOL ECON, 69:2060–2068. https://doi.org/10.1016/j.ecolecon.2010.06.010

Folkersen, M.V., 2018. Ecosystem valuation: Changing discourse in a time of climate change. ECOSYST SERV, 29:1–12. https://doi.org/10.1016/j.ecoser.2017.11.008

Fu, Y., Zhang, J., Zhang, C., Zang, W., Guo, W., Qian, Z., Liu, L., Zhao, J. and Feng, J., 2018. Payments for Ecosystem Services for watershed water resource allocations. J HYDROL, 556:689–700. https://doi.org/10.1016/j.jhydrol.2017.11.051

Geijzendorffer, I.R., Martin-Lopez, B. and Roche, P.K., 2015. Improving the identification of mismatches in ecosystemservices assessments. ECOL INDIC, 52:320–331. https://doi.org/10.1016/j.ecolind.2014.12.016

Grizzetti, B., Liquete, C., Antunes, P., Carvalho, L., Geamana, N., Giuca, R., Leone, M., McConnell, S., Preda, E., Santos, R., Turkelboom, F., Vadineanu, A. and Woods, H., 2016. Ecosystem services for water policy: Insights across Europe. ENVIRON SCI POLICY, 66:179–190. https://doi.org/10.1016/j.envsci.2016.09.006

Guo, Z.W., Xiao, X.M., Gan, Y.L. and Zheng, Y.J., 2001. Ecosystem functions, services and their values - a case study in Xingshan County of China. ECOL ECON, 38:141–154. https://doi.org/10.1016/S0921-8009(01)00154-9

Hackbart, V.C.S., de Lima, G.T.N.P. and Dos Santos, R.F., 2017. Theory and practice of water ecosystemservices valuation: Where are we going? ECOSYST SERV, 23:218–227. https://doi.org/10.1016/j.ecoser.2016.12.010

Hayha, T. and Franzese, P.P., 2014. Ecosystem services assessment: A review under an ecological-economic and systems perspective. ECOL MODEL, 289:124–132. https://doi.org/10.1016/j.ecolmodel.2014.07.002

Hayha, T., Franzese, P.P., Paletto, A. and Fath, B.D., 2015. Assessing, valuing, and mapping ecosystemservices in Alpine forests. ECOSYST SERV, 14:12–23. https://doi.org/10.1016/j.ecoser.2015.03.001

Kenter, J.O., Bryce, R., Christie, M., Cooper, N., Hockley, N., Irvine, K.N., Fazey, I., O’Brien, L., Orchard-Webb, J., Ravenscroft, N., Raymond, C.M., Reed, M.S., Tett, P. and Watson, V., 2016. Shared values and deliberative valuation: Future directions. ECOSYST SERV, 21:358–371. https://doi.org/10.1016/j.ecoser.2016.06.006

Kubiszewski, I., Costanza, R., Anderson, S. and Sutton, P., 2017. The future value of ecosystem services: Global scenarios and national implications. ECOSYST SERV, 26:289–301. https://doi.org/10.1016/j.ecoser.2017.05.004

Liu D., Hu Z.T., and Jin L.S., 2018. Review on analytical framework of eco-compensation. ACTA ECOLOGICA SINICA, 38: 380–392 (in Chinese with English abstract). https://doi.org/10.5846/stxb201605281028

Mancini, M.S., Galli, A., Coscieme, L., Niccolucci, V., Lin, D., Pulselli, F.M., Bastianoni, S. and Marchettini, N., 2018. Exploring ecosystemservices assessment through Ecological Footprint accounting. ECOSYST SERV, 30:228–235. https://doi.org/10.1016/j.ecoser.2018.01.010

Martin, D.M. and Mazzotta, M., 2018. Non-monetary valuation using Multi-Criteria Decision Analysis: Sensitivity of additive aggregation methods to scaling and compensation assumptions. ECOSYST SERV, 29:13–22. https://doi.org/10.1016/j.ecoser.2017.10.022

Millennium Ecosystem Assessment, 2005. Ecosystems and Human Well-Being: Synthesis. Island Press, Washington, DC.

Morri, E., Pruscini, F., Scolozzi, R. and Santolini, R., 2014. A forest ecosystemservices evaluation at the river basin scale: Supply and demand between coastal areas and upstream lands (Italy). ECOL INDIC, 37:210–219. https://doi.org/10.1016/j.ecolind.2013.08.016

Pagiola, S. and Platais, G., 2006. Payments for Environmental Services: From Theory to Practice. World Bank, Washington DC.

Pascual, U., Balvanera, P., Díaz, S., Pataki, G.R., Roth, E., Stenseke, M., Watson, R.T., Ba Ak Dessane, E., Islar, M., Kelemen, E., Maris, V., Quaas, M., Subramanian, S.M., Wittmer, H., Adlan, A., Ahn, S., Al-Hafedh, Y.S., Amankwah, E., Asah, S.T., Berry, P., Bilgin, A., Breslow, S.J., Bullock, C., Cáceres, D., Daly-Hassen, H., Figueroa, E., Golden, C.D., Gómez-Baggethun, E., González-Jiménez, D., Houdet, J.L., Keune, H., Kumar, R., Ma, K., May, P.H., Mead, A. O Farrell, P., Pandit, R., Pengue, W., Pichis-Madruga, R., Popa, F., Preston, S., Pacheco-Balanza, D., Saarikoski, H., Strassburg, B.B., van den Belt, M., Verma, M., Wickson, F. and Yagi, N., 2017. Valuing nature’s contributions to people: the IPBES approach. CURR OPIN ENV SUST, 26-27:7–16. https://doi.org/10.1016/j.cosust.2016.12.006

PRWRC: Pearl River Water Resources Commission of the Ministry of Water Resources, 2017. Zhujiang River Water Resources Bulletin 2016, Guangzhou (in Chinese). http://www.pearlwater.gov.cn/xxcx/szygg/16gb/

Schomers, S. and Matzdorf, B., 2013. Payments for ecosystemservices: A review and comparison of developing and industrialized countries. ECOSYST SERV, 6:16–30. https://doi.org/10.1016/j.ecoser.2013.01.002

SCPRC: State Council of the People’s Republic of China, 2014. The State Council approved the Development Planning of Zhujiang-Xijiang Economic Zone. State Council of the People’s Republic of China, Beijing (in Chinese). http://www.gov.cn/zhengce/content/2014-07/16/content_8933.htm

Sun, X., Crittenden, J.C., Li, F., Lu, Z. and Dou, X., 2018. Urban expansion simulation and the spatio-temporal changes of ecosystemservices, a case study in Atlanta Metropolitan area, USA. SCI TOTAL ENVIRON, 622-623:974–987. https://doi.org/10.1016/j.scitotenv.2017.12.062

TEEB Synthesis, 2010. Mainstreaming the Economics of Nature: a Synthesis of the Approach, Conclusions and Recommendations of TEEB. Earthscan, London and Washington.

Ulanowicz, R.E., 1986. Growth and Development: Ecosystems Phenomenology. Springer-Verlag, New York.

Wei, H., Fan, W., Wang, X., Lu, N., Dong, X., Zhao, Y., Ya, X. and Zhao, Y., 2017. Integrating supply and social demand in ecosystemservices assessment: A review. ECOSYST SERV, 25:15–27. https://doi.org/10.1016/j.ecoser.2017.03.017

Wunder, S., 2015. Revisiting the concept of payments for environmental services. ECOL ECON, 117:234–243. https://doi.org/10.1016/j.ecolecon.2014.08.016

Wunder, S., Engel, S. and Pagiola, S., 2008. Taking stock: A comparative analysis of payments for environmental services programs in developed and developing countries. ECOL ECON, 65:834–852. https://doi.org/10.1016/j.ecolecon.2008.03.010

Zheng, H., Robinson, B.E., Liang, Y., Polasky, S., Ma, D., Wang, F., Ruckelshaus, M., Ouyang, Z. and Daily, G.C., 2013. Benefits, costs, and livelihood implications of a regional payment for ecosystem service program. P NATL ACAD SCI USA, 110:16681–16686. https://doi.org/10.1073/pnas.1312324110

